# The microRNA, miR-133b, functions to slow Duchenne muscular dystrophy pathogenesis

**DOI:** 10.1101/2020.03.19.988519

**Authors:** Thomas Taetzsch, Dillon Shapiro, Randa Eldosougi, Tracey Myers, Robert Settlage, Gregorio Valdez

## Abstract

Duchenne muscular dystrophy (DMD) is characterized by progressive degeneration of skeletal muscles. To date, there are no treatments available to slow or prevent the disease. Hence, it remains essential to identify molecular factors that promote muscle biogenesis since they could serve as therapeutic targets for treating DMD. While the muscle enriched microRNA, miR-133b, has been implicated in the biogenesis of muscle fibers, its role in DMD remains unknown. To assess the role of miR-133b in DMD-affected skeletal muscles, we genetically ablated miR-133b in the *mdx* mouse model of DMD. In the absence of miR-133b, the tibialis anterior muscle of juvenile and adult *mdx* mice is populated by small muscle fibers with centralized nuclei, exhibits increased fibrosis, and thickened interstitial space. Additional analysis revealed that loss of miR-133b exacerbates DMD-pathogenesis partly by altering the number of satellite cells and levels of protein-encoding genes, including previously identified miR-133b targets as well as genes involved in cell proliferation and fibrosis. Altogether, our data demonstrate that skeletal muscles utilize miR-133b to mitigate the deleterious effects of DMD.

## Introduction

Duchenne muscular dystrophy (DMD) is the most prevalent of the muscular dystrophies, primarily affecting boys, with symptoms arising between 3-5 years of age. While advances in medical care have allowed DMD patients to live into adulthood, there is still no cure for the disease [1]. DMD is caused by loss of function of the dystrophin gene, an X-linked gene important for maintaining the integrity of the muscle fiber sarcolemma [1–4] through its association with dystroglycan and sarcoglycans [2,5,6]. While the initial formation of skeletal muscles proceeds mostly unimpeded during development, muscle fibers rapidly degenerate as individuals mature due to damages on the peripheral membrane caused by mechanical stress associated with muscle contraction [1,2]. As the disease becomes more severe, skeletal muscle progenitor cells, known as satellite cells, fail to adequately proliferate and differentiate in order to replace damaged muscle fibers [7]. DMD-pathogenesis is further exacerbated by chronic immune cell infiltration and fibrosis [8,9] that results from deposition of collagens and fat emanating from fibroblasts and adipocytes [7,10,11]. These pathological changes invariably impair the function and stability of remaining muscle fibers, compromising voluntary movements and ultimately causing death of affected individuals [1]. Thus, it is important to continue to explore new avenues for treating DMD.

The progressive nature of DMD [1] suggests that endogenous mechanisms could be recruited to mitigate damages caused by DMD and to maintain the regenerative capacity of skeletal muscles. Arguably, the most attractive endogenous molecular mechanisms would be those that bolster the regenerative capacity of skeletal muscles and endow muscle fibers with the ability to overcome fibrosis and inflammation resulting from the loss of dystrophin [12]. In this regard, microRNAs have received attention as candidate molecules for modifying DMD-pathology because of their functions in normal and disease-affected skeletal muscles and the relative ease with which they can be packaged into vectors [13–16]. These ubiquitous small non-coding RNAs are approximately 22 nucleotides in length and impact a wide range of cellular processes by inhibiting translation of target mRNAs [17,18].

There is ample evidence suggesting that the muscle-enriched microRNA, miR-133b, may be a candidate for modifying DMD-pathogenesis. In vitro studies have shown that targeting miR-133b and its close relative, miR-133a, affect different stages of myogenesis, including the proliferation of satellite cells [19,20] and myoblasts [21], the differentiation of satellite cells into either myoblasts or brown adipose tissue [22], and the formation of muscle fibers [20,23,24]. In vivo, published data indicate that skeletal muscles upregulate miR-133b during regeneration following injury [25] and may utilize miR-133b to maintain homeostasis during cellular, functional and biochemical changes induced by exercise [26] and denervation [27] as well as in sarcopenia [28]. Additionally, miR-133b was shown to accelerate muscle regeneration following injury in young adult rats alongside miR-1 and miR-206 [29]. Importantly, there is evidence indicating that miR-133 is elevated in DMD patients, however this study did not distinguish miR-133b from miR-133a [30]. Despite these findings, the role of miR-133b in DMD remains unclear. This is because miR-133b has yet to be examined independently of other non-coding RNAs in DMD [31,32]. In one published study, miR-133b was proposed to collaborate with a long noncoding RNA, linc-MD1, to affect the timing and rate of muscle fiber formation in DMD [31]. In another study, it was shown that simultaneously deleting miR-133b and miR-206, another muscle enriched microRNA found in the same long non-coding RNA as miR-133b, has no effect on DMD pathogenesis in dystrophin-null (*mdx)* mice [32]. Therefore, it remains unknown if and how miR-133b plays a role in DMD.

To directly test the role of miR-133b in DMD-pathogenesis, we generated *mdx* mice lacking miR-133b. Analysis of these mice during various stages of disease severity revealed that miR-133b does in fact play important roles in DMD. Loss of miR-133b decreased the size of muscle fibers while increasing fibrosis in *mdx* mice. These changes are partly due to alterations in satellite cell abundance and changes in expression of a myriad of genes, including both known targets of miR-133b and genes shown to be involved in cell cycle regulation and promotion of fibrosis. Altogether, these findings strongly suggest that skeletal muscles increase miR-133b levels to mitigate damages caused by DMD.

## Results

### miR-133b is induced in dystrophin-deficient mice

Although miR-133 is elevated in the serum of DMD patients [30], it remains unknown if this is due to increased expression of miR-133b, miR-133a, or both. Additionally, the temporal relationship between miR-133b levels and severity of DMD in skeletal muscles remains unknown. To answer these questions, we examined miR-133b expression during the initial stage, postnatal day 30 (p30), and severe stage (p60), of the disease in the tibialis anterior (TA) muscle of *mdx* mice [33]. Since one nucleotide distinguishes mature miR-133b from miR-133a, we examined levels of pre-miRNAs instead of mature miRNAs, using qPCR. This analysis revealed that miR-133b is increased at p30 and p60 in *mdx* mice compared to control mice (Fig. 1A). Interestingly, miR-133b expression in *mdx* mice is significantly lower at p60 compared to p30. Thus, miR-133b levels are inversely correlated with the severity of the disease; highest at p30 and lowest at p60. This expression pattern was specific to miR-133b compared to miR-133a, which was expressed at similar or lower levels in *mdx* muscle (Fig. 1B). It also differed from other muscle-enriched microRNAs in *mdx* mice. While miR-206 was significantly higher at both ages in *mdx* mice, it is expressed at higher levels at p60 compared to p30 in *mdx* mice (Fig. s1A) unlike miR-133b, which is lower at p60 compared to p30 in *mdx* mice. Levels of miR-1 were similar at p30 and somewhat decreased at p60 in the TA muscle of *mdx* mice compared to control mice (Fig. s1B), confirming previous reports [34].

**Figure 1.**
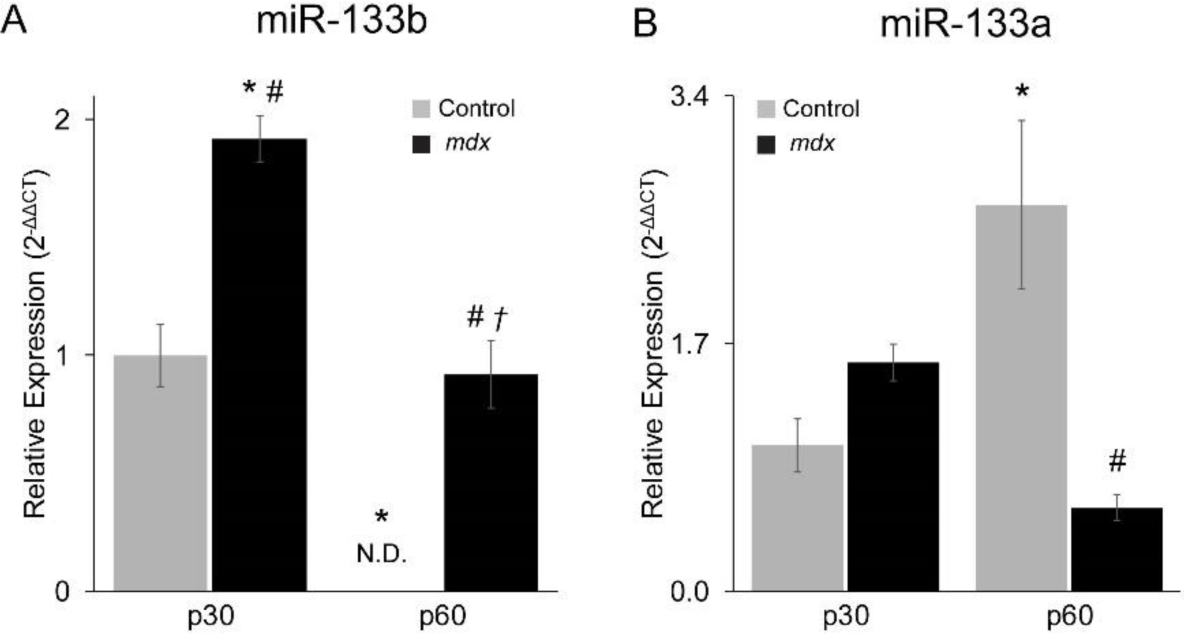
Analysis of miR-133b and miR-133a levels in *mdx* mice. Levels of precursor (A) miR-133b and (B) miR-133a in the TA muscle of juvenile (p30) and adult (p60) control and *mdx* mice were assessed using qPCR. All values reported as mean ± SEM; p30, n = 4; p60 n = 3. * p < 0.05 versus p30 control, # p < 0.05 versus p60 control, † p < 0.05 versus p30 *mdx*. N.D. not detected.

### Deletion of miR-133b exacerbates DMD-pathogenesis in skeletal muscles of *mdx* mice

To date, the role of miR-133b in DMD pathogenesis remains debated mainly because it has yet to be examined independently of the precursor long-noncoding RNA in which both miR-133b and miR-206 are encoded (LINCMD1) [16,35]. Previous work has demonstrated directly opposing myogenic functions of miR-206, miR-133b and other regions of the LINCMD1 gene [31]. While deletion of miR-206 alone exacerbates DMD-related pathology [25], the simultaneous deletion of mir-133b and miR-206 has no effect on it [32]. To examine the function of miR-133b independently of miR-206 in DMD, we used mice lacking only miR-133b (miR-133b^-/-^). First, we examined muscles only lacking miR-133b in otherwise healthy young adult mice. We found that deletion of miR-133b does not cause obvious changes in skeletal muscles. There is no difference in the average muscle fiber cross-sectional area (CSA, Fig s2 A,B,D), distribution of muscle fiber size (Fig. s2 C), mononucleated cells occupying the interstitial space (Fig. s2 E), and number of Pax7^+^ satellite cells (Fig. s2 F) between miR-133b^+/+^ and miR-133b^-/-^ mice. In stark contrast, deletion of miR-133b from *mdx* mice, following crossing of miR-133b null with *mdx* mice (herein referred to as *mdx*; miR-133b^-/-^), exacerbated DMD-related pathological features in skeletal muscles. Analysis of muscle fibers in the TA muscle of juvenile (p30) mice revealed a decrease in the muscle fiber CSA in *mdx*; miR-133b^-/-^ mice as compared to *mdx*; miR-133b^+/+^ (Figure 2 A,B,D). This reduction in muscle fiber CSA is due to fewer muscle fibers with large CSAs and more muscle fibers with small CSAs (Figure 2 A-C). As expected, the CSA of the entire TA muscle also decreased in *mdx*; miR-133b^-/-^ mice compared to *mdx;* miR-133b^+/+^ mice (Figure 2 E). Interestingly, the number of muscle fibers with centralized nuclei, a characteristic of regenerating juvenile and adult muscle fibers, was lower in *mdx*; miR-133b^-/-^ muscle however this difference was not statistically significant (Figure 2 F). Altogether, these data suggest that miR-133b affects the initial progression of DMD by altering the maturation and not the rate of muscle fiber degeneration and regeneration in *mdx* mice.

**Figure 2.**
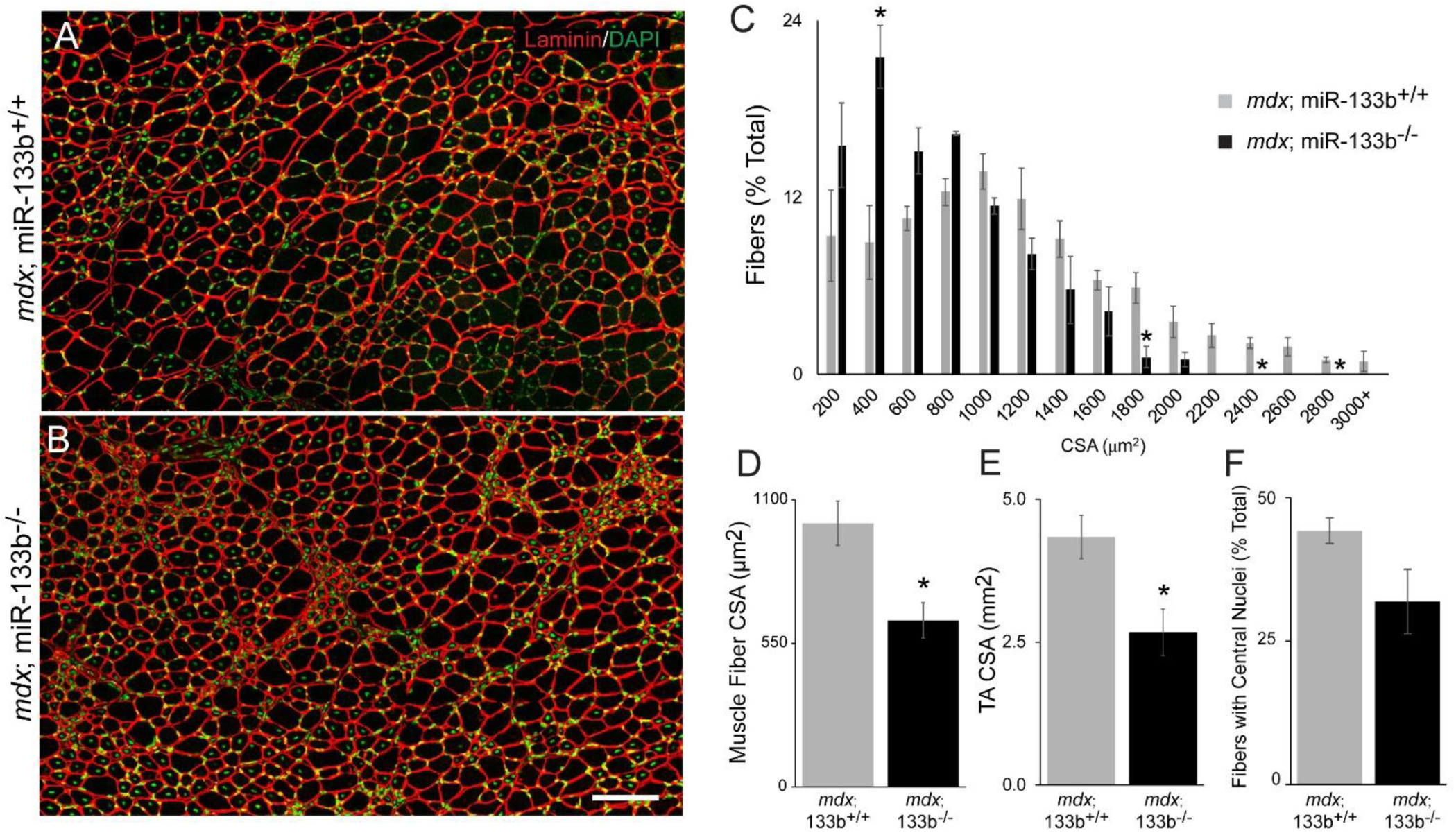
miR-133b plays an important role in slowing DMD pathogenesis in juvenile (p30) *mdx* mice. (A-B) Representative images of laminin (red) and DAPI (green) staining in TA muscle cross-sections used to determine muscle fiber cross-sectional area (CSA) in *mdx*; miR-133b^+/+^ and *mdx*; miR-133b^-/-^ mice. (C) A frequency distribution graph of muscle fiber CSAs shows decreased numbers of muscle fibers with larger CSAs in *mdx*; miR-133b^-/-^ mice. (D) The average muscle fiber CSA and (E) the average CSA of the TA muscle are significantly smaller in *mdx*; miR-133b^-/-^ as compared to *mdx*; miR-133b^+/+^ mice. (F) Regenerating muscle fibers, characterized by the presence of centralized nuclei, in TA muscle. All values reported as mean ± SEM, n = 3. * p < 0.05, scale bar = 100 µm.

We next assessed the effect of deleting miR-133b in *mdx* mice during stages of severe (p60) and plateauing (p90) DMD pathology [33]. At P60, the average muscle fiber and overall TA CSA was similar between *mdx* littermates with and without miR-133b (Figure 3 A,B,D,E). However, a frequency distribution analysis revealed significantly more muscle fibers with very small CSAs in the TA muscle of *mdx*; miR-133b^-/-^ mice (Figure 3 C). Additionally, the incidence of muscle fibers with centralized nuclei was significantly higher in *mdx*; miR-133b^-/-^ mice (Figure 3F). Thus, the continued presence of small muscle fibers, along with an increased number of regenerating fibers, demonstrates that loss of miR-133b also exacerbates DMD-related pathology at p60, when the disease is most severe in *mdx* mice. These differences between *mdx*; miR-133b^-/-^ and *mdx*; miR-133b^+/+^ muscle are mostly missing at p90, an age when muscle fibers have partly adapted to loss of dystrophin and exhibit less DMD-related pathology [33]. While the number of regenerating fibers remains slightly elevated in *mdx*; miR-133b^-/-^ muscle (Figure s3 F, p = 0.06), the average muscle fiber CSA (Figure s3 A-D) and overall size of the TA muscle (Figure s3 E) were similar between *mdx*; miR-133b^-/-^ and *mdx*;miR-133b^+/+^ littermates at p90.

**Figure 3.**
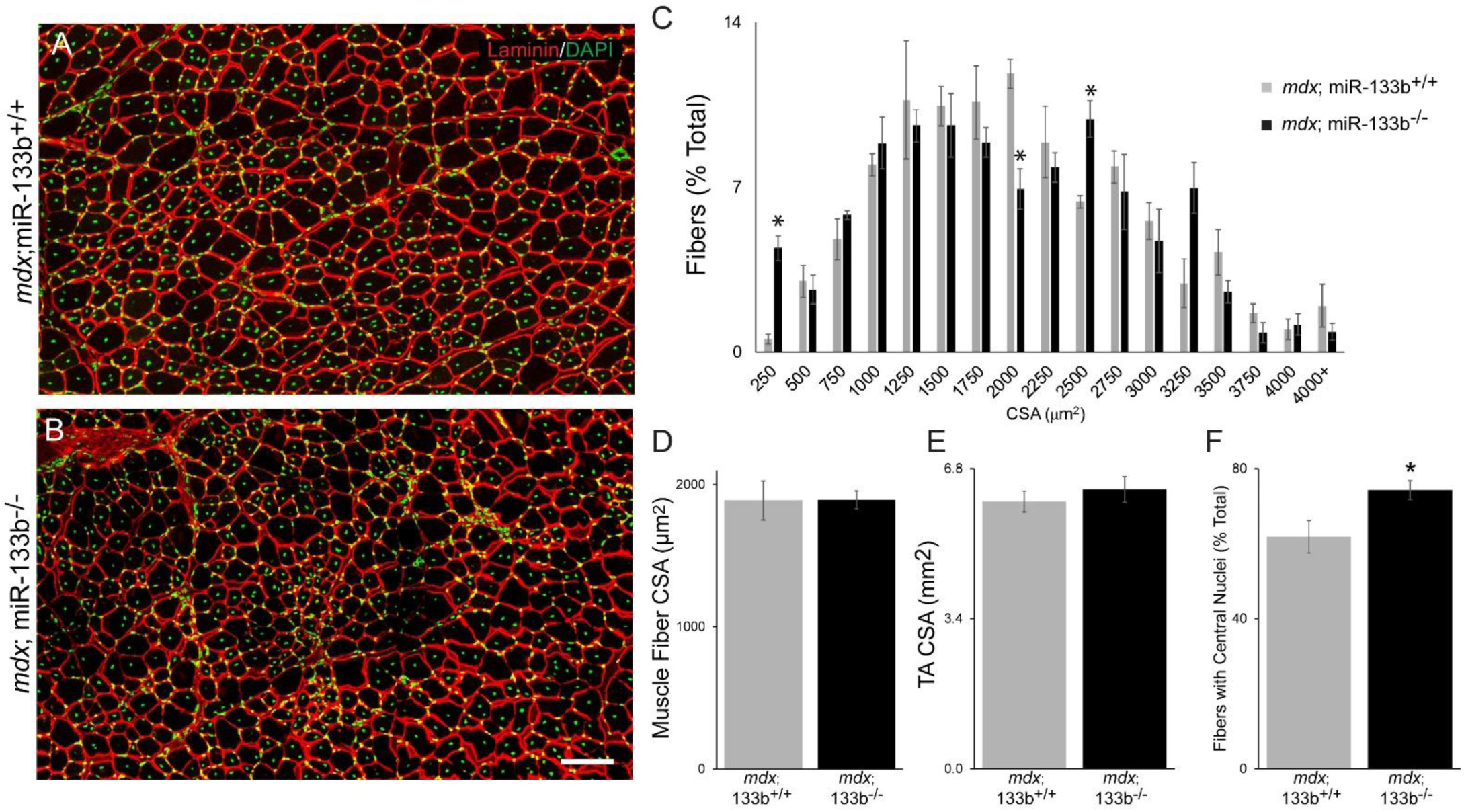
The loss of miR-133b moderately affects DMD pathogenesis in p60 *mdx* mice. Representative images of laminin (red) and DAPI (green) staining of p60 (A) *mdx*; miR-133b^+/+^ and (B) *mdx*; miR-133b^-/-^ TA cross sections reveals (C) increased numbers of muscle fibers with the smallest CSAs in *mdx*; miR-133b^-/-^ as compared to *mdx*; miR-133b^+/+^ muscle. (D) The average muscle fiber CSA and (E) the overall TA muscle size are similar between genotypes. (F) An increased number of regenerating muscle fibers, characterized by the presence of centralized nuclei, was observed in *mdx*; miR-133b^-/-^ TA muscle. All values reported as mean ± SEM, n = 3. * p < 0.05, scale bar = 100 µm.

Although we previously showed that neuromuscular junctions (NMJs) are unaffected in miR-133b null mice [27], it is possible that NMJs afflicted with DMD require miR-133b. To address this possibility, we examined NMJs in the extensor digitorum longus (EDL) muscle of *mdx*; miR-133b^-/-^ mice following staining for nicotinic acetylcholine receptors (nAChRs) with fluorescently-tagged alpha-bungarotoxin (fBTX). This analysis showed that loss of miR-133b resulted in diminished nAChR area (Fig. s4 A,B,D) with a similar trend in fragmentation (Fig. s4 C) and endplate area (Fig. s4 E) in p30 *mdx*; miR-133b^-/-^ versus *mdx*; miR-133b^+/+^ EDL. Despite these differences, the number of NMJs was similar between genotypes indicating that loss of miR-133b affects the morphology and not the number of muscle fibers in p30 *mdx* mice. In contrast, NMJs were indistinguishable at p60 between *mdx*;miR-133b^-/-^ mice and *mdx*;miR-133b^+/+^ mice (Fig. s4 G-L). This finding is in line with the muted effect of miR-133b deletion on skeletal muscle of *mdx* mice at p60.

### Loss of miR-133b increases fibrosis and presence of infiltrating cells in *mdx* muscles

Fibrosis and inflammation are prominent features of DMD pathology, and contribute to progressive loss of myofibers and muscle weakness [9,10]. We thus asked if loss of miR-133b exacerbates DMD by increasing fibrosis and the number of cells occupying the interstitial space of skeletal muscle. We measured the area of the interstitial space in the TA muscle of p30 *mdx*; miR-133b^+/+^ and *mdx*; miR-133b^-/-^ mice by tracing the laminin-positive region separating muscle fibers. This analysis showed a marked increase in the interstitial space in *mdx*; miR-133b^-/-^ compared to *mdx*; miR-133b^+/+^ mice (Fig. 4A-C). We then counted the number of nuclei residing in the interstitial space as a rough measurement of infiltrating cells. We found that the number of nuclei occupying the interstices of myofibers is more than two-fold higher in the TA muscle of *mdx*; miR-133b^-/-^ compared to *mdx*; miR-133b^+/+^ mice (Fig. 4D). We extended these analyses to P60 mice and found a similar trend with a roughly two-fold increase in the area of laminin positive staining (Fig. 4E-G) and twice as many nuclei (Fig. 4H) in P60 *mdx*; miR-133b^-/-^ compared to *mdx; miR-133b*^*+/+*^littermates. These results show that deletion of miR-133b exacerbates DMD-pathogenesis partly by increasing fibrosis and the number of mononucleated cells in the interstitial space, two cellular changes shown to be associated with diminished health of muscle fibers. The increased presence of mononucleated cells in the interstitial space may be indicative of an elevated presence of inflammatory cells, which typically infiltrate DMD muscle tissue [9]. Once again, we found that deleting miR-133b at p90 did not further exacerbate fibrosis and accumulation of mononucleated cells in the interstitial space (Fig. 4 I-L). These findings indicate that other mechanisms override the effect of deleting miR-133b and the loss of dystrophin around p90 in *mdx* mice.

**Figure 4.**
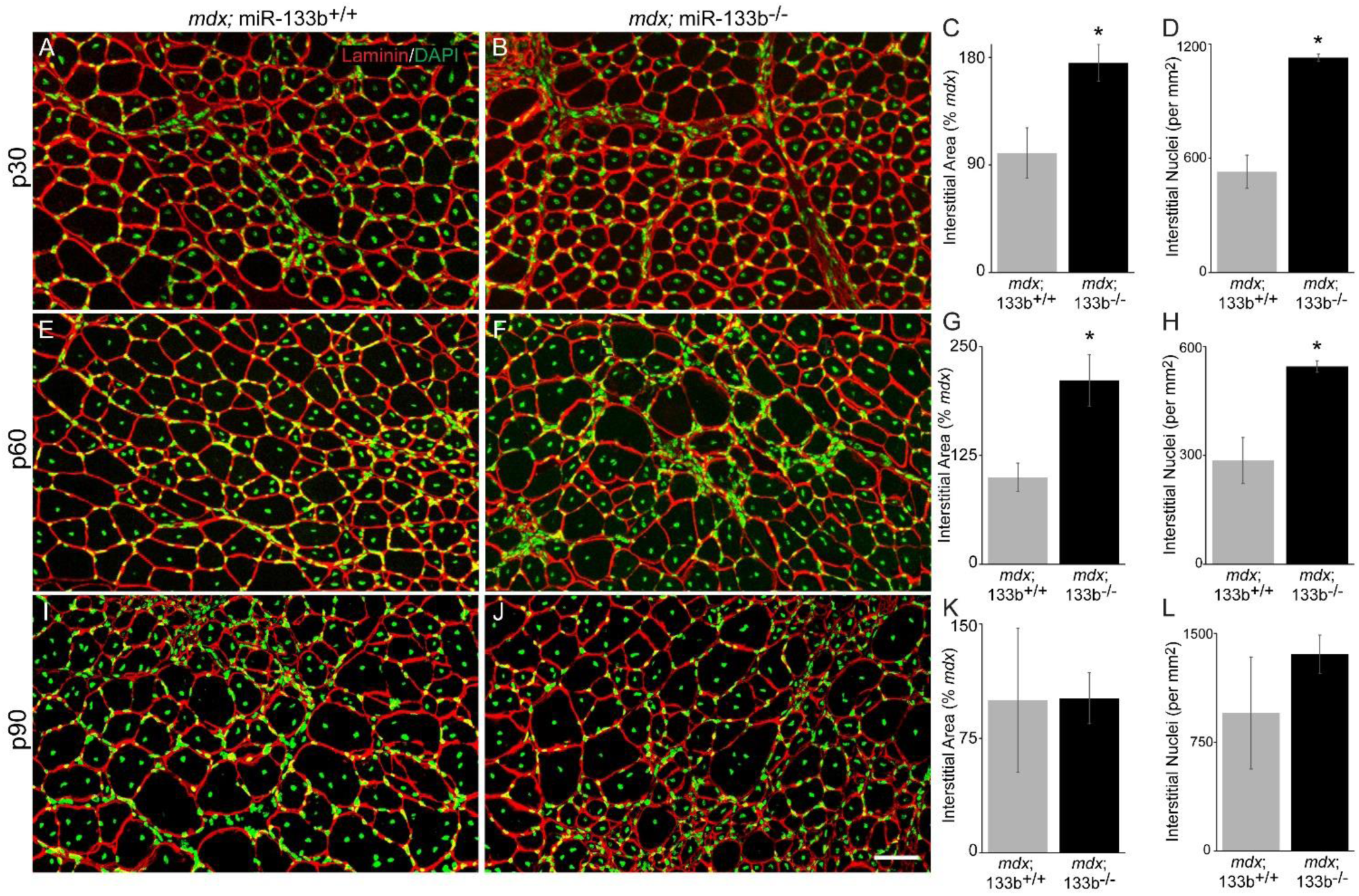
Fibrosis and cell infiltration are increased in *mdx*; miR-133b^-/-^ mice. (A-B) Laminin (red) and DAPI (green) staining in cross-sections of the TA muscle was used to assess the area of the interstitial space as a proxy for fibrosis in *mdx*; miR-133b^+/+^ and *mdx*; miR-133b^-/-^ mice. (C) Increased laminin^+^ interstitial area was observed in p30 *mdx*; miR-133b^-/-^ versus *mdx*; miR-133b^+/+^ TA muscle. (D) DAPI^+^ nuclei within the interstitial space were analyzed as a measurement of infiltrating cells. Increased numbers of nuclei within the interstitial space were observed in p30 *mdx*; miR-133b^-/-^ TA muscle. (E-G) At p60, a similar trend of increased laminin^+^ interstitial area and (H) increased interstitial nuclei was observed in *mdx*; miR-133b^-/-^ TA muscle. (I-K) At p90 interstitial area and (L) interstitial nuclei were similar in *mdx*; miR-133b^-/-^ and *mdx*; miR-133b^+/+^ TA muscle. All values reported as mean ± SEM, n = 3. * p < 0.05, scale bar = 50 µm.

### Impact of miR-133b deletion on satellite cells in *mdx* mice

In vitro studies have uncovered important roles for miR-133b in satellite cell proliferation and differentiation [14,36], raising the possibility that loss of miR-133b exacerbates DMD-pathogenesis by affecting satellite cells. To address this possibility, we examined the number of Pax7^+^ satellite cells in the TA muscle of p30, p60 and p90 *mdx*; miR-133b^+/+^ and *mdx*; miR-133b^- /-^ mice. At p30, the number of Pax7^+^ satellite cells was slightly elevated in *mdx*; miR-133b^-/-^ as compared to *mdx*; miR-133b^+/+^ muscle (Fig. 5 A,C,D). However as the disease progressed, the number of Pax7^+^ satellite cells decreased at a faster rate in *mdx*; miR-133b^-/-^ compared to *mdx*; miR-133b^+/+^ muscle at both p60 (Fig. 5 B,C,E) and p90 (Fig. 5C and s5). These results show that miR-133b affects DMD-pathogenesis by altering the number of satellite cells.

**Figure 5.**
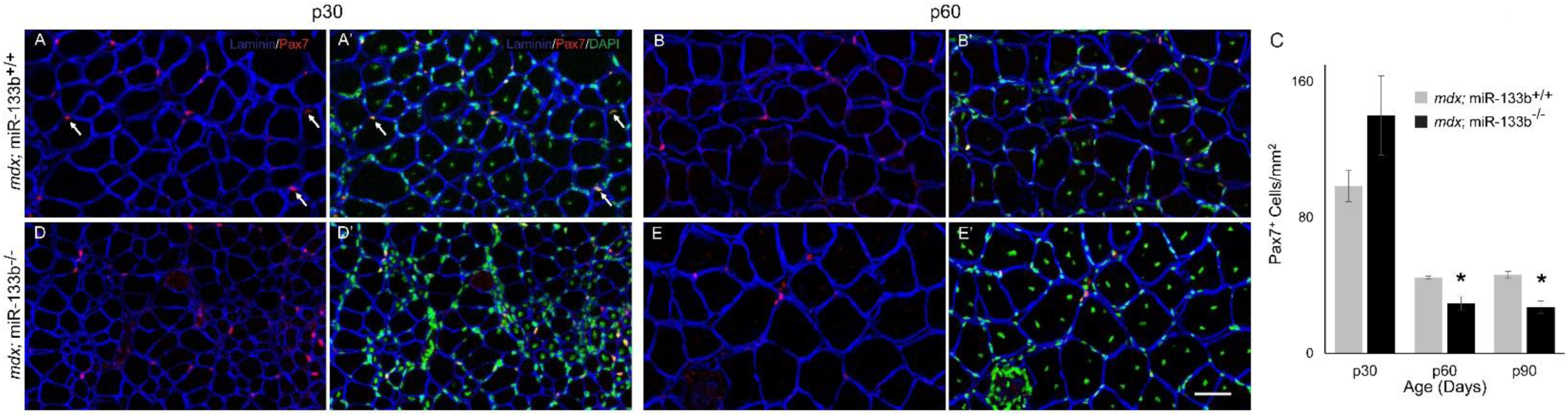
The number of satellite cells is altered in *mdx*; miR-133b^-/-^ muscle. (A,B,D,E) Representative images of laminin (blue), Pax7 (red) and DAPI (green) immunohistochemistry performed on TA cross sections collected from p30 and p60 *mdx*; miR-133b^+/+^ and *mdx*; miR-133b^-/-^ mice. White arrows in panel A indicate examples of Pax7/DAPI double positive nuclei used to identify satellite cells. (C) Quantification of satellite cell density shows a slight elevation in the number of Pax7^+^ satellite cells at p30 and decreased numbers of Pax7^+^ satellite cells at p60 and p90 in *mdx*; miR-133b^-/-^ as compared to *mdx*; miR-133b^+/+^ muscle. All values reported as mean ± SEM. p30 and p90, n = 4; p60, n = 3. * p < 0.05, scale bar = 50 µm.

### Deletion of miR-133b moderately affects muscle regeneration in acutely stressed skeletal muscles in healthy young adult mice

The results above, showing that deletion of miR-133b exacerbates DMD-pathogenesis, raised the prospect that miR-133b also plays a critical role in the repair of skeletal muscles affected by other stressors. We thus asked if miR-133b must be present for muscle fibers to regenerate properly and in a timely fashion following an acute injury. For this, we administered cardiotoxin (CTX), which causes muscle fibers to rapidly degenerate [37,38], to the TA muscle of miR-133b^+/+^ and miR-133b^-/-^ littermates. As muscle fibers begin to reform, around 7 days (7d) post-CTX injection, the average muscle fiber CSA was lower and there were fewer muscle fibers with larger CSAs in miR-133b^-/-^ as compared to miR-133b^+/+^ mice (Fig. 6 A,B,E). By 14d post-CTX injection, there was a slight increase in the number of fibers with very small CSAs (<200 µm^2^) and fewer muscle fibers with CSAs that are typical of fully regenerated muscle (i.e. >2200 um^2^) in miR-133b^-/-^ as compared to miR-133b^+/+^ muscle (Fig. 6 C,D,F). These changes led to a net reduction in the average muscle fiber CSA in miR-133b^-/-^ mice (Fig. 6 G) and demonstrate that regenerated muscle fibers mature at a slower rate in the absence of miR-133b. These differences do not appear to result from changes in the interstitial space since the number of nuclei was similar in miR-133b^-/-^ versus miR-133b^+/+^ muscle at both 7d and 14d post-CTX (Fig. 6H). The slower rate of muscle fiber regeneration also does not seem to be caused by a lower abundance of Pax7^+^ satellite cells (Fig. 6I and Fig. s6). Thus, miR-133b does not have an obvious impact on satellite cells and other mononucleated cells in the interstitial space in transiently stressed skeletal muscles. This is in stark contrast to chronically stressed skeletal muscles in *mdx* mice, where miR-133b must be present to minimize the accumulation of pathological features.

**Figure 6.**
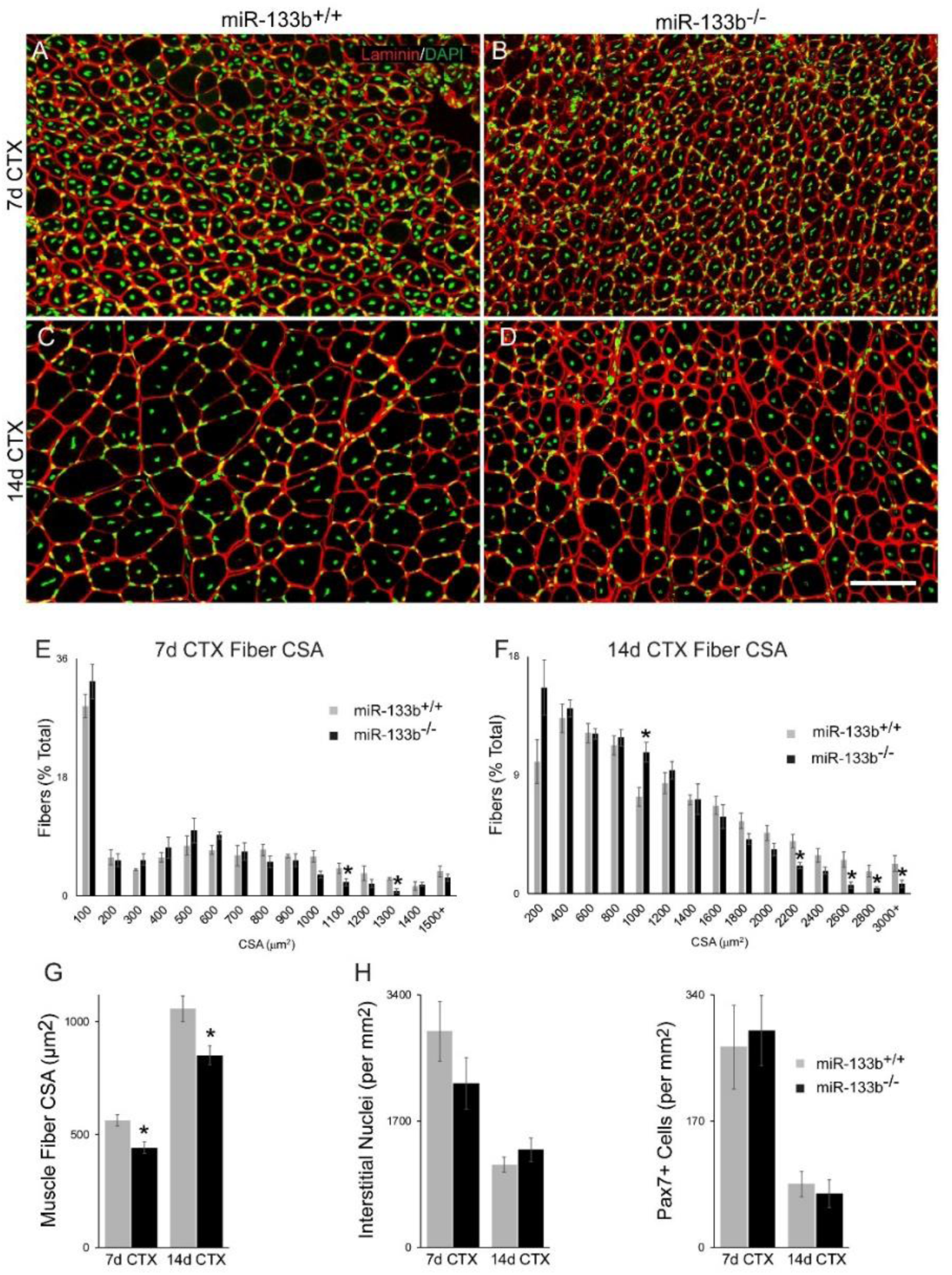
Deletion of miR-133b from control mice moderately delays muscle fiber regeneration following an acute injury. Cardiotoxin (CTX) was administered to the TA muscle of healthy p60 miR-133b^+/+^ and miR-133b^-/-^ littermates to induce muscle regeneration. (A-D) Representative images of laminin (red) and DAPI (green) staining in muscle collected at 7d (A,B) and 14d (C,D) post-CTX from miR-133b^+/+^ and miR-133b^-/-^ muscle. (E) A frequency distribution graph of muscle fiber CSAs at 7d post-CTX shows fewer muscle fibers with larger CSAs in miR-133b^-/-^ versus miR-133b^+/+^ TA. (F) At 14d post-CTX a similar trend of fewer muscle fibers with larger CSAs was observed in miR-133b^- /-^ muscle. (G) At both 7 and 14d post-CTX administration a decrease in average muscle fiber CSA was observed in miR-133b^-/-^ muscle. (H) Numbers of nuclei in the interstitial space and (I) numbers of Pax7^+^ satellite cells were similar in miR-133b^-/-^ versus miR-133b^+/+^ TA at both 7d and 14d post-CTX. All values reported as mean ± SEM. 7d CTX, n= 4; 14d CTX, n = 6. * p < 0.05, scale bar = 50 µm.

### Deleting miR-133b accelerates satellite cell proliferation but slows the formation of myotubes

To determine if loss of miR-133b directly impacts satellite cell proliferation and their ability to generate new myofibers, we isolated and cultured satellite cells from p15-p20 miR-133b^+/+^ and miR-133b^-/-^ littermates at low density. After 72 h *in vitro*, Pax7^+^ satellite cells lacking miR-133b proliferated faster compared to miR-133b^+/+^ controls (Fig. 7 A-C). To determine whether miR-133b impacts satellite cell differentiation and fusion into myotubes, we seeded satellite cells at a high density and treated with differentiation media. At both 2 (Fig. 7 D,E) and 5 (Fig. 7 F,G) days post-differentiation we observed smaller myosin^+^ myotubes in miR-133b^-/-^ versus miR-133b^+/+^ cultures (Fig. 7 H), suggesting an impairment of myotube formation in the absence of miR-133b. Taken together, these results show that miR-133b functions to control satellite cell proliferation and formation of myofibers, which are two cellular processes altered in in *mdx*; miR-133b^-/-^ muscle.

**Figure 7.**
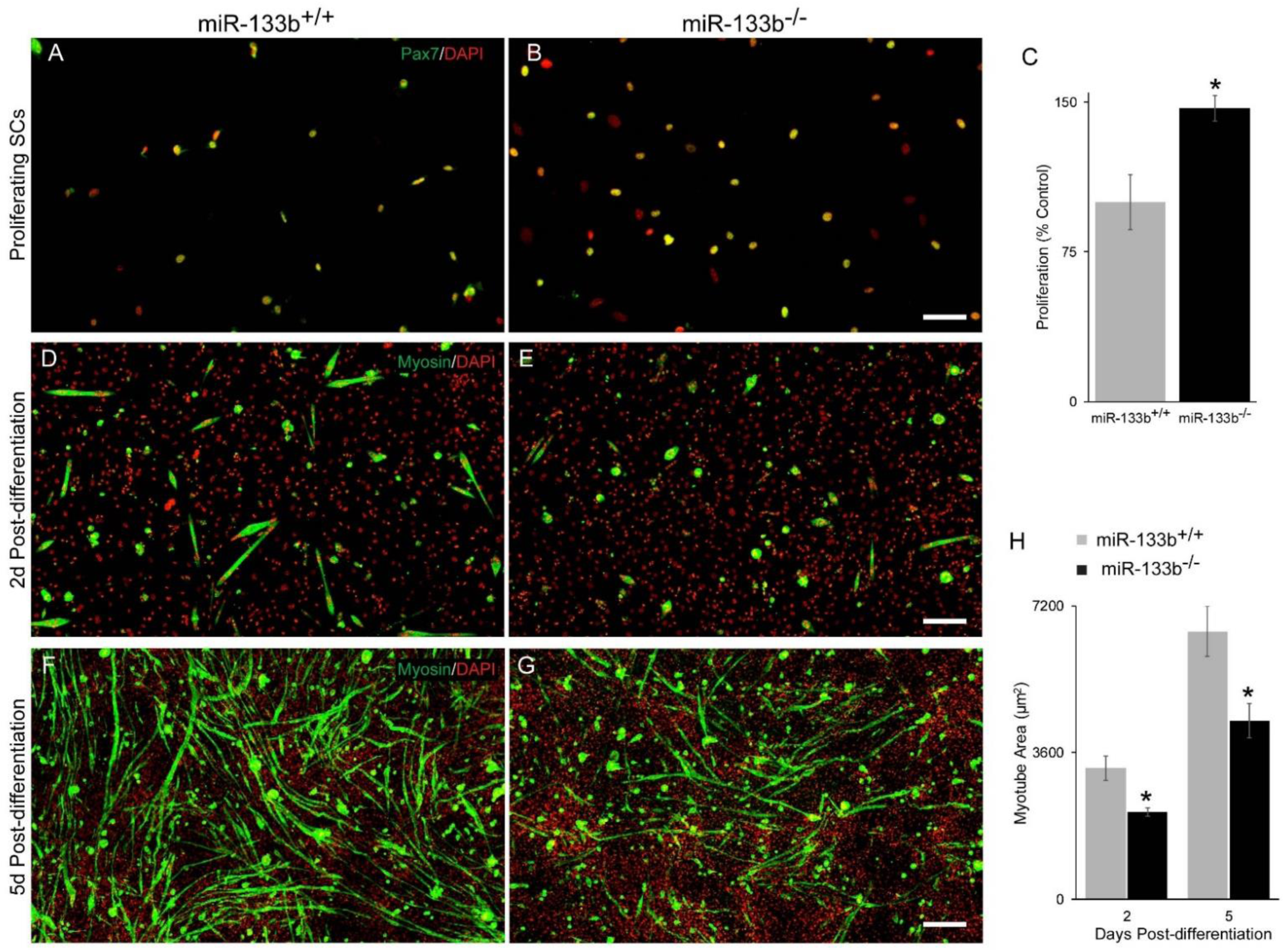
Deletion of miR-133b enhances satellite cell proliferation but inhibits myotube formation. (A,B) Representative images of Pax7 (green) and DAPI (red) immunocytochemistry of miR-133b^+/+^ and miR-133b^-/-^ primary satellite cell-enriched cultures at 72 h *in vitro*. (C) Satellite cell proliferation was determined by measuring the ratio of Pax7^+^ satellite cells at 72 h versus <16 h *in vitro.* Enhanced rates of proliferation of Pax7^+^ satellite cells were observed in miR-133b^-/-^ as compared to miR-133b^+/+^ primary satellite cell-enriched cultures. n = 4. (D,E) Representative images of myosin (green) and DAPI (red) immunocytochemistry at 2d post-differentiation (PD) in miR-133b^+/+^ and miR-133b^-/-^ primary satellite cell-enriched cultures. (F,G) Myosin/DAPI immunocytochemistry at 5d PD. (H) Analysis of average myotube area shows that myotubes are smaller in miR-133b^-/-^ versus miR-133b^+/+^ primary satellite cell-enriched cultures at 2d and 5d PD. 2d PD, n =3; 5d PD, n = 4. All values reported as mean ± SEM, * p < 0.05, scale bar = 200 µm.

### Molecular signatures associated with loss of miR-133b in *mdx* mice

The increased DMD-pathogenesis in *mdx*; miR-133b^-/-^ mice indicates that miR-133b impacts expression of protein-coding genes with important functions in DMD-afflicted skeletal muscles. To identify genes altered in the TA muscle lacking miR-133b, we compared the transcriptome between wild type, miR-133b^-/-^, *mdx*; miR-133b^+/+^, and *mdx*; miR-133b^-/-^ mice using RNA seq. Genes with an absolute log fold change greater than 1 and adjusted p-value less than 0.01 were deemed differentially expressed between genotypes. This analysis revealed the following: 1) there are very few genes differentially expressed between miR-133b^+/+^ and miR-133b^-/-^ mice (Fig. 8A), partly corroborating cellular analysis showing that loss of miR-133b has no obvious effect on otherwise healthy skeletal muscles (Fig. s2) and NMJs [27]; 2) a myriad of genes are significantly altered in the TA muscle of *mdx* compared to wild type control mice (Fig. 8 B); 3) deletion of miR-133b further alters the transcriptome of the TA muscle in *mdx* mice (Fig. 8C).

**Figure 8.**
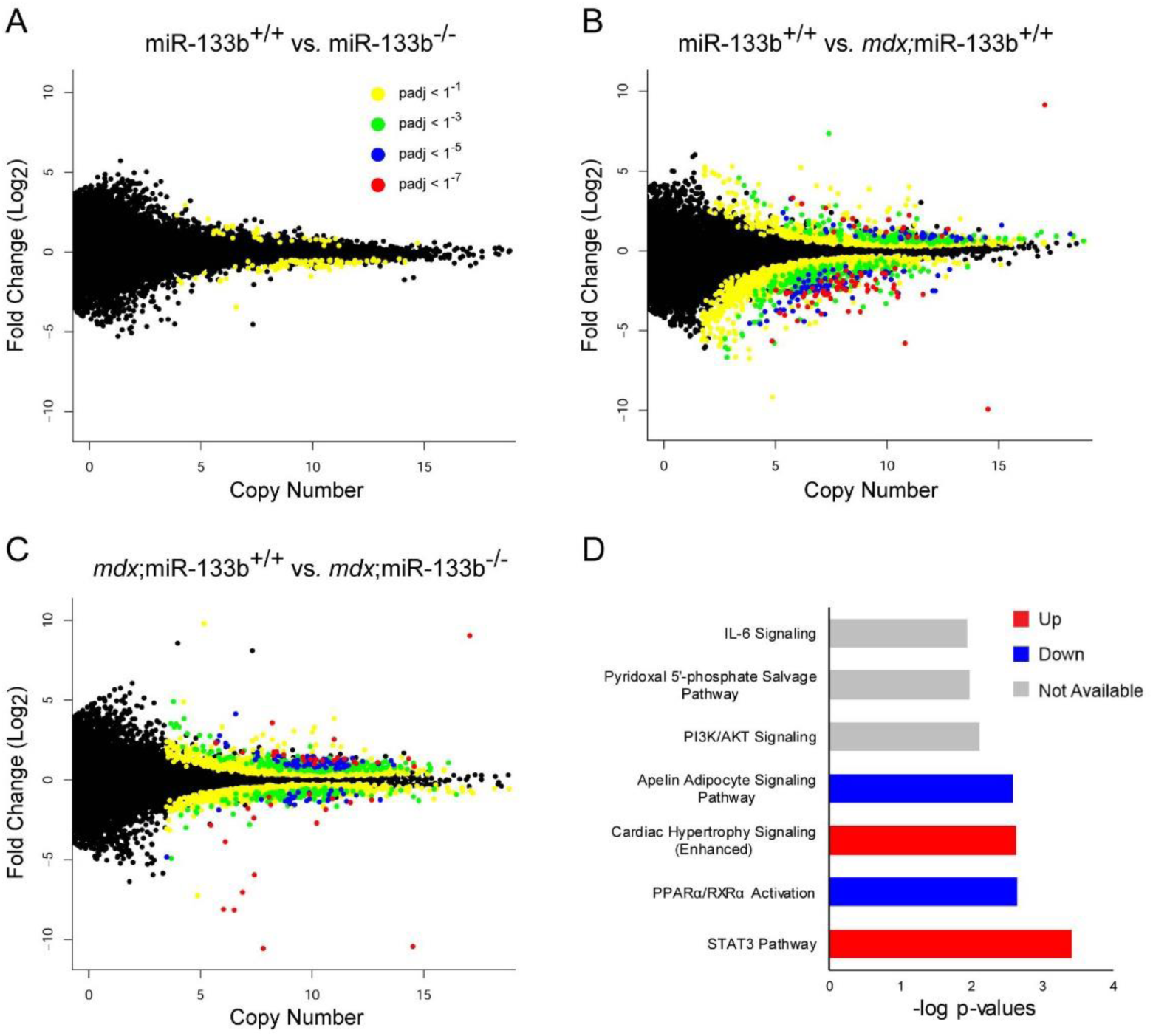
Transcriptomic changes resulting from miR-133b deletion in healthy and *mdx* juvenile (p30) TA muscle. (A) Scatter plot comparison of differentially expressed genes in miR-133b^+/+^ versus miR-133b^-/-^ TA muscle, as identified by RNA seq. Copy number in miR-133b^+/+^ muscle is represented on the x-axis and log2 fold change versus miR-133b^-/-^ is represented on the y-axis. Adjusted p-value for each gene expression comparison is represented by dot coloration. (B) Scatter plot comparison of genes expressed in healthy (miR-133b^+/+^) versus DMD-afflicted (*mdx*; miR-133b^+/+^) muscle. Copy number in miR-133b^+/+^ muscle is represented on the x-axis and log2 fold change versus *mdx*; miR-133b^+/+^ is represented on the y-axis, p-values color coded according to legend in panel A. (C) Scatter plot comparison of genes expressed in *mdx*; miR-133b^+/+^ versus *mdx*; miR-133b^-/-^ muscle. Copy number in *mdx*; miR-133b^+/+^ muscle is represented on the x-axis and log_2_ fold change versus *mdx*; miR-133b^-/-^ muscle is represented on the y-axis, p-values color coded according to legend in panel A. (D) Ingenuity Pathway Analysis of genes with a fold change absolute value of greater than 2 and adjusted p-value of less than 0.01 in *mdx*; miR-133b^+/+^ versus *mdx*; miR-133b^-/-^ muscle identifies the top signaling pathways impacted by deletion of miR-133b. n = 4.

To gain insights into the signaling pathways miR-133b alters to exacerbate DMD-pathology in *mdx* mice, we used Ingenuity Pathway Analysis (IPA). This revealed several pathways with particular relevance to DMD pathogenesis that were closely associated with deletion of miR-133b in *mdx* mice (Fig. 8D) including STAT3, Apelin, and IL-6. STAT3 has been shown to be expressed by satellite cells where it plays a pivotal role in myogenesis by promoting satellite cell proliferation and inhibiting the formation of myotubes [39]. Recent studies have demonstrated that. IL-6 is closely associated with infiltrating cells of the interstitial space and is downregulated in response to glucocorticoid treatment in DMD patients [39]. Apelin has been characterized as an exercised-induced signaling molecule that prevents sarcopenia, improves muscle regeneration, and decreases inflammation in aging muscle [40]. Modulation of these pathways by miR-133b may serve to protect muscle from DMD pathogenesis.

Prominent among genes upregulated in *mdx*; miR-133b^-/-^ muscle were a number of known miR-133b targets that were identified using miRTarBase [41] that have also been shown to be involved in TGF-B signaling and DMD pathology (Table 1). These included RhoA, a well characterized target of miR-133b [42] that is associated with myoblast fusion, muscle regeneration, fatty infiltration and effectiveness of glucocorticoid treatment in *mdx* mice [43–47]. A number of other genes related to TGF-β signaling that are not presently known to be direct miR-133b targets were also upregulated in *mdx*; miR-133b^-/-^ muscle, including the TGF-β receptor, TGF-β-induced factor homeobox 1 (TGIF1), SMAD3, and SMAD5 (Table 2) [48,49]. In addition, latent TGF-β binding protein 4 (LTBP4), a protein that is responsible for sequestering TGF-β in the extracellular matrix, and one of a handful of genes known to affect disease progression in DMD patients [50,51], was downregulated in *mdx*; miR-133b^-/-^ muscle. Overall, this expression dataset supports important roles for miR-133b in mitigating DMD pathogenesis. It also provides promising candidates, in the form of specific genes and signaling modules, that could be directly targeted to attenuate the various pathological features associated with DMD.

## Discussion

This is the first study to directly assess the role of miR-133b on DMD pathogenesis and on the regenerative capacity of skeletal muscle in vivo. We show that deletion of miR-133b reduces the size of muscle fibers and increases fibrosis in *mdx* mice. Importantly, we provide cellular and molecular mechanistic insights on how miR-133b impacts DMD pathogenesis. We demonstrate that deletion of miR-133b alters the number of satellite cells and their ability to form myofibers. We also uncover a myriad of genes and signaling modules that are altered when miR-133b is deleted in *mdx* mice. Altogether, our findings provide the basis to use miR-133b and downstream targets to augment the regenerative capacity of skeletal muscles possibly through satellite cells, and to reduce fibrosis and infiltration of mononucleated cells to treat DMD.

The impact of miR-133b on satellite cells is a significant finding in this study because of the role these cells play in the progression of DMD pathogenesis. Specifically, it is clear that satellite cell dysfunction contributes to the loss of skeletal muscle mass and increased presence of fibrotic and adipose tissue that accompanies the progression of DMD [11,12]. We show that miR-133b plays an important role in modulating the number of satellite cells and their capacity to generate myofibers in DMD. Without miR-133b, satellite cells were more numerous during the initial stages of DMD but then significantly decreased with disease progression in *mdx* mice. These changes in satellite cell numbers were accompanied by an increase in the number of small myofibers during the early and severe stages of the disease in *mdx* mice. In culture, we found that miR-133b deletion resulted in elevated satellite cell proliferation and inhibited myotube formation. These in vitro findings suggest that a central role of miR-133b in satellite cells is to decrease the duration of satellite cell activation by promoting not only differentiation but also quiescence. Hence, one possible explanation for the initial increase and subsequent decrease in satellite cell numbers at p30 and p60, respectively, in *mdx*;miR-133b^-/-^ mice is that deletion of miR-133b results in depletion of the quiescent satellite cell pool that is necessary to maintain the satellite cell population as the disease progresses. Thus, in the absence of miR-133b, there are initially more satellite cells at the onset of the disease as satellite cells favor proliferation over both quiescence and differentiation. However, over time, a preference for activation ultimately depletes the quiescent satellite cell pool which in turn causes an overall decrease in newly generated activated satellite cells and an insufficient regenerative response to muscle fiber damage. Irrespective, these data add to published studies indicating that miR-133b plays important roles in satellite cell proliferation and commitment to myoblasts. Thus, the varying levels of miR-133b at specific stages of DMD severity in *mdx* muscle (i.e. p30 and p60) may be necessary to both control the proliferation and differentiation of satellite cells to promote regeneration of muscle fibers.

In contrast to *mdx* mice, deletion of miR-133b from healthy control mice had a limited effect on skeletal muscles. Muscle fiber size, satellite cell number, ECM thickness, and levels of infiltrating cells were normal in healthy muscle lacking miR-133b. While over 600 genes were differentially regulated in *mdx* muscle in the absence of miR-133b, RNA seq analysis identified only two differentially regulated genes related to deletion of miR-133b in healthy muscle. These findings are in line with previous studies demonstrating that miRNAs are primarily utilized by cells in mature tissues to coordinate responses under stress [14,26,52,53]. It also supports our findings that miR-133b is utilized by activated satellite cells, which are limited in number in healthy skeletal muscle. In addition to impacting DMD pathogenesis, we found that deletion of miR-133b slows the rate of muscle fiber regeneration following cardiotoxin-induced injury. However, it must be noted that the number of satellite cells was similar between miR-133b^+/+^ and miR-133b^-/-^ skeletal muscles following cardiotoxin-induced injury. This could be due to the earliest time point at which we examined satellite cells (7d post-CTX), which does not represent the peak of proliferation [54]. Alternatively, it may indicate that miR-133b plays a role in ensuring the continuous proliferation of satellite cells during chronic, asynchronous muscle regeneration and is not necessary for repair of transiently stressed muscle fibers.

In addition to orchestrating muscle regeneration, this study demonstrates an additional and previously unknown role for miR-133b in mitigating fibrosis in DMD muscle. Global deletion of miR-133b enhanced the severity of DMD-associated fibrosis, as seen by increased interstitial space and infiltration of cells within it. While elevated fibrosis may be directly related to dysregulated satellite cell proliferation and impaired myofiber formation, it is possible that miR-133b may directly regulate fibroblasts, adipocytes, immune cells, and other cells residing in skeletal muscles. Several lines of evidence support roles for miR-133b in other types of muscle resident cells. Previous work has demonstrated that miR-133 blocks adipogenesis in skeletal muscle tissue [22]. Our RNA seq study identified alterations in the expression of a number of genes related to inflammation and fibrosis in *mdx*; miR-133b-/- muscle, including TGF-B pathway members such as SMAD3, LTBP4, and TGFBR. In addition, IPA identified cell signaling associated with the inflammatory cytokine IL-6 to be prominently affected by deletion of miR-133b from *mdx* muscle. The therapeutic relevance of IL-6 has been demonstrated by a recent study showing that a reduction in IL-6 is associated with improved outcomes associated with steroid treatment in DMD patients [39].

Loss of satellite cells, inability to reform muscle fibers, fibrosis, and accumulation of mononucleated cells in the interstitial space are hallmarks of DMD pathology. These cellular and molecular features inevitably contribute to progressive loss of muscle fibers, muscle weakness, and death. Hence, it is critical to continue to identify targets that could slow these features of DMD. Our results provide evidence that miR-133b, a short oligonucleotide, could be used to mitigate DMD pathology by augmenting the proliferation and differentiation of satellite cells while reducing inflammation and fibrosis. We also identified a myriad of candidate genes that may also be attractive therapeutic targets in DMD. Thus, future experiments will assess the impact of manipulating miR-133b levels and the action of downstream targets at specific stages of DMD pathology.

## Methods

### Animals

Dystrophin-null DMD mice (*mdx*, Jackson Labs 001801 [55]) were purchased from Jackson Laboratories (Bar Harbor, Maine). miR-133b^-/-^ mice were obtained from the Feng lab [56]. Wild-type control mice used for qPCR were on a mixed genetic background and were age and sex-matched to *mdx* mice. miR-133b^-/-^ mice were crossed with *mdx* mice to generate *mdx*; miR-133b^-/-^ mice on a mixed genetic background. Male mice were used for all comparisons of *mdx*; miR- 133b^+/+^ with *mdx*; miR-133b^-/-^. Male and female mice were used for assessing deletion of miR-133b in healthy muscle (i.e. miR-133b^+/+^ vs miR-133b^-/-^). Breeding, housing and experimental use of animals were performed in accordance with the National Institutes of Health and Virginia Tech Institutional Animal Care and Use Committee guidelines.

### Immunohistochemistry (IHC): Whole mounted muscle

Mice were anesthetized with isoflurane and perfused transcardially with phosphate buffered saline followed by 4% paraformaldehyde. Following dissection, the extensor digitorum longus (EDL) muscle was incubated with Alexafluor 555 conjugated alpha-bungarotoxin (#B35451, Invitrogen, Carlsbad, CA) diluted 1:1000 in PBS for 2 h at room temperature, washed 3 times with PBS for 5 minutes and mounted to a glass slide in Vectashield (Vector Laboratories, Burlingame, CA).

### IHC: Muscle Cross Sections

The tibialis anterior (TA) muscle was dissected from paraformaldehyde perfused mice, post-fixed in 30% sucrose in PBS for 48 h at 4C, cross-sectioned at 16 µm in Tissue Freezing Medium (#TFM-5, General Data, Cincinnati, OH) with a cryostat, and mounted to gelatin coated glass slides. For laminin IHC, TA sections were incubated in blocking buffer (5% BSA, 3% goat serum, 0.1% Triton X-100 in PBS) for 1 h at room temperature, incubated in primary antibody (Rabbit anti-laminin, #L9393, Sigma-Aldrich, St. Louis, MO) diluted 1:250 in blocking buffer overnight at 4C, washed 3 times in PBS, incubated in Alexafluor 568 conjugated anti-rabbit antibody (#A11036, Invitrogen, Carlsbad, CA) diluted 1:1000 in blocking buffer for 1 h at room temperature, washed twice in PBS, incubated in DAPI (#D1306, Fisher Scientific, Waltham, MA) diluted 1:1000 in PBS for 20 minutes at room temperature, and washed twice in PBS before Vectashield (Vector Laboratories) mounting medium and coverslip application. Pax7 IHC was performed according to [57], briefly, sections were incubated in M.O.M. blocking buffer (#MKB-2213, Vector Laboratories, Burlingame, CA) for 1 h at room temperature, incubated in mouse anti-Pax7 antibody (undiluted, Antibody Registry ID: AB_528428) and rabbit anti-laminin (1:250, #L9393,Sigma-Aldrich, St. Louis, MO) overnight at 4°C, washed in PBS 3 times for 5 minutes, incubated in Alexafluor 546 conjugated anti-mouse antibody (#A-11010, Invitrogen, Carlsbad, CA) diluted 1:1000 in 5% horse serum for 1 h at room temperature, washed twice in PBS, incubated in DAPI (1:1000 in PBS) for 20 minutes at room temperature, and washed twice in PBS before Vectashield mounting medium and coverslip application. Pax7 antibody was a generous donation from Dr. Julia von Maltzahn.

### Immunocytochemistry (ICC)

Cell cultures were fixed in 4% paraformaldehyde for 30 minutes at room temperature and washed 3 times in PBS for 5 minutes. Pax7 immunocytochemistry was performed according to Pax7 IHC protocol above. Myosin ICC was performed with mouse anti-myosin (1:100, #MF20, DSHB, Iowa City, IA) according to laminin IHC protocol above.

### Imaging

Imaging was performed with a Zeiss LSM 710 laser scanning confocal microscope (Carl Zeiss Microscopy, Berlin, Germany) using a 20× (0.8 numerical aperture) objective. Zeiss Zen software was used for maximum intensity projections and stitching of tile scans.

### Muscle fiber cross sectional area and central nuclei analyses

Muscle fiber analysis was performed on four tile scan images of laminin stained TA cross sections. Cross sections of 16 µm thickness were selected from the medial TA at 0.3 mm intervals. Muscle fibers were identified by laminin positive rings within muscle tissue. Muscle fibers with central nuclei were defined by the presence of a DAPI-positive nucleus within laminin positive rings that are not immediately adjacent to the laminin positive staining. The fractionator method [58] was used to randomly sample an average of 290 muscle fibers per muscle. To achieve this, a grid was superimposed on the images with ImageJ software and all muscle fibers located on a grid point were selected for measurement. Muscle fiber CSA measurements were performed with ImageJ. Interstitial Nuclei Analysis Interstitial nuclei counts were performed on laminin/DAPI stained 16 µm TA cross sections using the fractionator sampling method. On average, 30 regions measuring 7800 µm^2^ were randomly sampled across two cross sections collected from the medial TA. Interstitial nuclei were identified as DAPI positive nuclei located in areas of thickened laminin deposition in the interstitial space and not immediately adjacent to muscle fibers. Counting and sampling were performed on blinded images with ImageJ software.

### Interstitial Space Analysis

Fibrosis analysis was performed on laminin/DAPI stained 16 µm thick TA cross sections using the fractionator sampling method. On average, 22 regions measuring 20,000 µm^2^ were randomly sampled across two cross sections collected from the medial TA. Interstitial area was identified by the presence of thick laminin staining occupied by nuclei in the interstices of muscle fibers. Sampling and measurements were obtained from blinded images with ImageJ software and reported as a percent of the total cross-sectional area sampled.

### *In Vivo* Satellite Cell Counts

Counting of Pax7+ satellite cells was performed on blinded images obtained from Pax7/laminin/DAPI stained TA cross sections of 16 µm thickness using the fractionator sampling method. Four cross sections were selected from the medial TA at 0.3 mm intervals. Pax7+ cells were identified by the presence of a Pax7+/DAPI+ nucleus located immediately adjacent to a laminin+ muscle fiber.

### Satellite Cell Proliferation Analysis

Satellite cell proliferation analysis was performed on blinded images of satellite cell enriched cultures using the fractionator sampling method. Following muscle dissociation from a single mouse, cells were seeded to two chamber slides. The first slide was fixed immediately after the cells attached to the slide (4-16 h post-seed) and the second slide was fixed at 72 h post-seed. The cell cultures were stained with Pax7/DAPI, imaged, and satellite cells were identified by the presence of a Pax7+/DAPI+ nucleus. One replicate represents 3 technical replicates derived from the dissociation of skeletal muscles of a single miR-133b^+/+^ mouse or its miR-133b^-/-^ littermate. Data are presented as the average fold change in the number of Pax7+ cells at 72 h post-seed relative to the number of Pax7+ cells immediately after seeding for a given replicate.

### Myotube Size

Myotube size analysis was performed on blinded images of primary muscle cultures derived from miR-133b^+/+^ and miR-133b^-/-^ littermates that were fixed at 2 or 5 d post-differentiation and stained with myosin/DAPI. Myotubes were identified by the presence of myosin and at least two nuclei. Area measurements were obtained by outlining the myosin^+^ area of a myotube using ImageJ software.

### Cardiotoxin Injury

Cardiotoxin from Naja melanoleuca (#v9000, Sigma-Aldrich, St. Louis, MO) was diluted in sterile PBS to a final concentration of 70 µg/mL. Following general anesthesia of p60 miR-133b^+/+^ and miR-133b^-/-^ littermates with ketamine (50 mg/kg IP), 50 µL of cardiotoxin solution was delivered to the belly of the TA muscle with a 31 gauge needle.

### Primary Myoblast Culture

Hindlimb muscles were collected from p15-p21 miR-133b^+/+^ and miR-133b^-/-^ littermates. After removal of connective tissue and fat, muscles were cut into 5 mm^2^ pieces with forceps and digested in 2 mg/mL collagenase II (Worthington Chemicals, Lakewood, NJ) for 1h at 37°C. Muscles were further dissociated by mechanical trituration in high glucose Dulbecco’s modified eagle medium (DMEM, #11965118, Life Technologies, Carlsbad, CA) containing 10% horse serum (Life Technologies #16050122) and passed through a 40 µm filter to generate a single cell suspension. Excess debris was removed from the suspension by centrifugation in 4% BSA followed by a second centrifugation in 40% Optiprep solution (Sigma-Aldrich, St. Louis, MO) from which the interphase was collected. Cells were washed in 10% horse serum in DMEM and seeded to laminin (#23017015, Life Technologies, Carlsbad, CA) coated chamber slides (NUNC Permanox, #160005, NUNC, Rochester, NY) in growth media consisting of 20% fetal bovine serum (#FP-0500-A, Atlas Biologicals, Fort Collins, CO), 0.1% penicillin/streptomycin (Life Technologies, Carlsbad, CA), 2 mm glutamine (Life Technologies, Carlsbad, CA), and 0.1% Fungizone (Life Technologies, Carlsbad, CA) in high glucose DMEM. For analysis of satellite cell proliferation, cells were seeded at a density of 20,000 cells per well. For analysis of myotube formation, cells were seeded at a density of 100,000 cells per well. Once cells were 100% confluent, growth medium was replaced with differentiation medium consisting of 2% horse serum (Life Technologies, Carlsbad, CA) and 0.1% penicillin/streptomycin in high glucose DMEM to induce myotube formation. Myotube cultures were fixed at 2 and 5 d after addition of fusion media. Cell cultures were maintained at 37°C and 5% CO_2_.

### mRNA analysis

RNA was extracted from the TA muscle using Trizol reagent (Life Technologies, Carlsbad, CA) and the Aurum Total RNA Mini Kit (Bio-Rad, Hercules, CA). Reverse transcription of RNA was performed with iScript™ Reverse Transcription Supermix (Bio-Rad, Hercules, CA). qPCR was performed with iTAQ SYBR Green Supermix (Bio-Rad, Hercules, CA) using the CFX Connect Real Time PCR System (Bio-Rad, Hercules, CA). Primers were used at a concentration of 300 nM, primer sequences used in this study are listed in Table 3. RNA sequencing was performed by the Genomics Sequencing Center at Virginia Tech using an mRNA-stranded Seq library prep (Illumina, San Diego, CA) and NextSeq High Output 75SR sequencing (Illumina, San Diego, CA). Following sequencing, data were trimmed for both adaptor and quality using a combination of ea-utils and Btrim [59,60]. Sequencing reads were aligned to the genome using Tophat2/HiSat [61] and counted via HTSeq [62]. QC summary statistics were examined to identify any problematic samples (e.g. total read counts, quality and base composition profiles (+/- trimming), raw fastq formatted data files, aligned files (bam and text file containing sample alignment statistics), and count files (HTSeq text files). Following successful alignment, mRNA differential expression was determined using contrasts of and tested for significance using the Benjamini-Hochberg corrected Wald Test in the R-package DESeq [63]. Signaling pathway analysis was generated through the use of IPA (QIAGEN Inc., https://www.qiagenbio-informatics.com/products/ingenuity-pathway-analysis).

### Statistical analysis

Independent Welch’s or Student’s T-tests were used to make comparisons between 2 means. One-way ANOVA with Bonferroni post-hoc analysis was used to make comparisons between 3 or more means. Statistical analyses performed with the Microsoft Excel Data Analysis plugin and R software. Data are expressed as mean ± SEM. A value of P < 0.05 was considered statistically significant.

## Acknowledgements

The authors wish to thank Dr. Julia von Maltzahn for donating the Pax7 antibody used in this study and for assistance with establishing a Pax7 IHC protocol for the laboratory.

