## Supplementary material for "The microRNA, miR-133b, functions to slow Duchenne muscular dystrophy pathogenesis": tables and supplementary figures

**Table 1.** miR-133b targets upregulated in *mdx*; miR-133b$^{-/-}$ versus *mdx*; miR-133b$^{+/+}$ TA muscle. Gene upregulation determined from RNA seq. Targets of miR-133b identified by the miRTarBase database.

| Gene | Function | Implications in DMD | Validation Methods | References |
|---|---|---|---|---|
| RhoA | GTPase | Impairs myoblast fusion and promotes adipogenesis | 4 | [43–47] |
| SP1 | Transcription factor | Regulates myogenesis | 2 | [64] |
| Tnrc6b | RNA-induced silencing | Myo-miR related myogenesis | 1 | [14,17] |
| PIAS2 | E3 ubiquitin ligase | Inhibits the satellite cell differentiation molecule STAT2 | 1 | [65] |
| PTBP2 | miRNA biogenesis | Myo-miR related myogenesis | 3 | [14,66] |

**Table 2.** TGF-β pathway components differentially regulated in *mdx*; miR-133b$^{-/-}$ as compared to *mdx*; miR-133b$^{+/+}$ TA muscle, as identified by RNA seq.

| **Gene** | **Name** | **Function** | **Up or down in absence of miR-133b?** | **References** |
|---|---|---|---|---|
| LTBP4 | Latent TGF-β binding protein 4 | Sequesters TGF-β in ECM | Down | [48] |
| SMAD3 | SMAD3 | Transcription factor | Up | [48] |
| SMAD5 | SMAD5 | Transcription factor | Up | [48] |
| TAB3 | TAK3/MAP3K7 binding protein 3 | Activate TGFB activated kinase | Up | [67] |
| TGFβR | TGFβ Receptor | Transmembrane receptor | Up | [48] |
| TGIF1 | TGFβ-induced factor homeobox 1 | Transcriptional co-repressor | Up | [49] |

**Table 3.** qPCR primers.

| Gene | Forward Primer (5'-3') | Reverse Primer (5'-3') |
|---|---|---|
| pre miR-1a | AGCACATACTTCTTTATGTACCCA | ACTTCTTTACATTCCATAGCACTGA |
| pre miR-133a | GGTAAAATGGAACCAAATCGCCT | GGTTGAAGGGGACCAAATCCA |
| pre miR-133b | GCTGGTCAAACGGAACCAAG | ATATTGAGCTTTGCCAGCCCT |
| pre miR-206 | GGCCACATGCTTCTTTATATCC | AAACCACACACTTCCTTACATTCC |

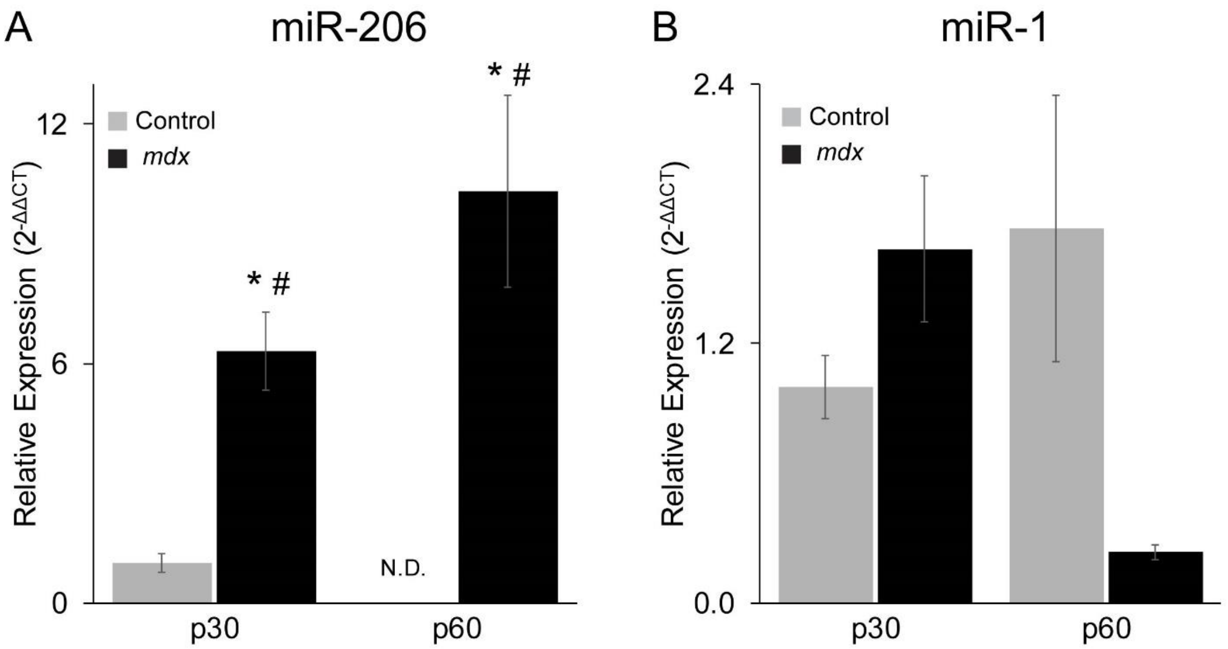


**Supplementary Figure 1.** Levels of muscle specific precursor (A) miR-206 and (B) miR-1 in juvenile (p30) and adult (p60) *mdx* and control TA muscle were determined by qPCR. All values reported as mean ± SEM; p30, n = 4; p60 n = 3. * $p < 0.05$ versus p30 control, # $p < 0.05$ versus p60 control. N.D. not detected.

*Fig. 2 Supplement 1*

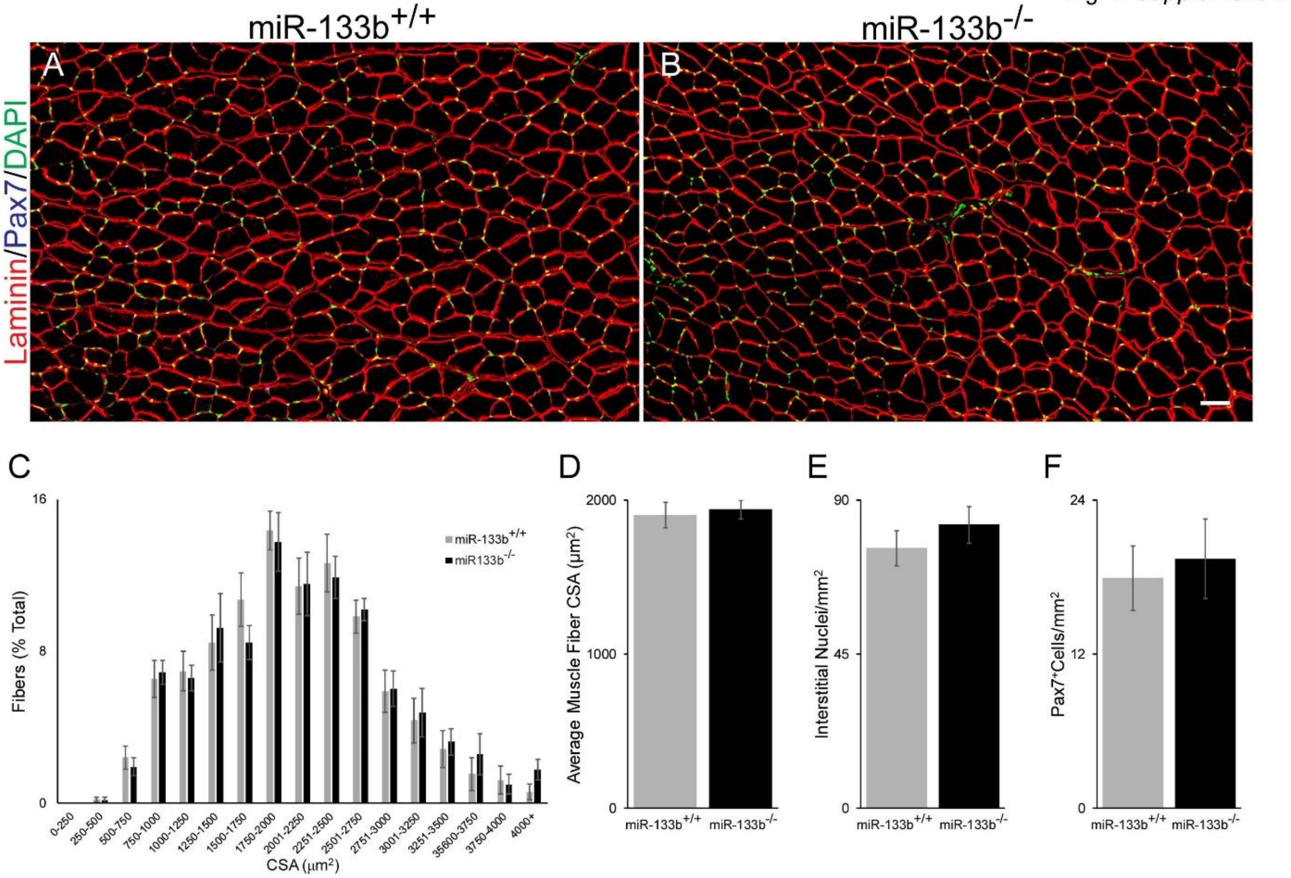


**Supplementary Figure 2. Analysis of muscle fibers in control mice lacking miR-133b.** (A,B) Pax7/laminin/DAPI IHC of p60 miR-133b$^{+/+}$ and miR-133b$^{-/-}$ TA cross sections shows no differences in (C) muscle fiber CSA distribution, (D) average muscle fiber CSA, (E) number of nuclei within the interstitial space, or (F) number of Pax7$^{+}$ satellite cells. miR-133b$^{+/+}$, n = 5; miR-133b$^{-/-}$, n = 4. All values reported as mean ± SEM; Scale bar = 50 μm.

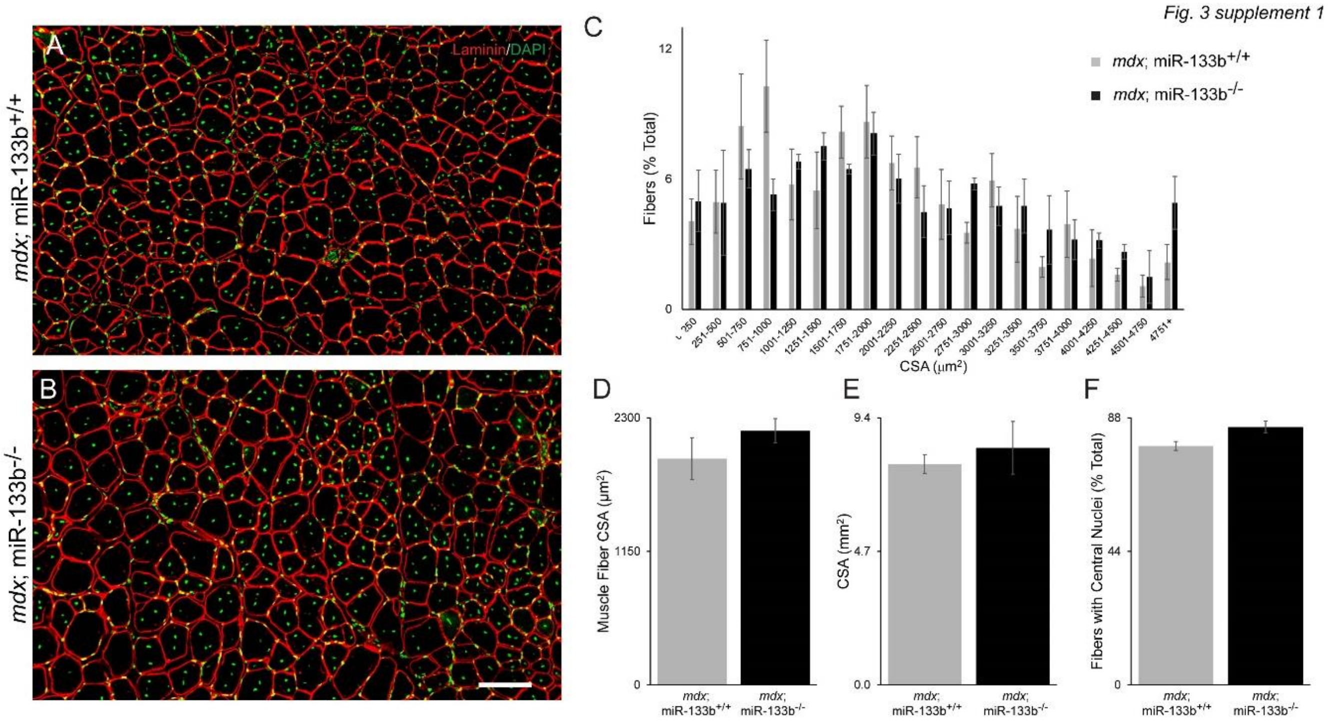


**Supplementary Figure 3.** Loss of miR-133b does not affect DMD pathogenesis in p90 *mdx* mice. (A,B) Laminin IHC of *mdx*; miR-133b$^{+/+}$ and *mdx*; miR-133b$^{-/-}$ TA cross sections shows (C) a similar distribution of muscle fiber sizes, (D) similar average muscle fiber size, and (E) similar overall TA size between genotypes. (F) A small increase in the number of regenerating muscle fibers, characterized by the presence of centralized nuclei, was observed in *mdx*; miR-133b$^{-/-}$ TA muscle, however this difference was not statistically significant. All values reported as mean ± SEM, n = 3. * $p < 0.05$, scale bar = 100 μm.

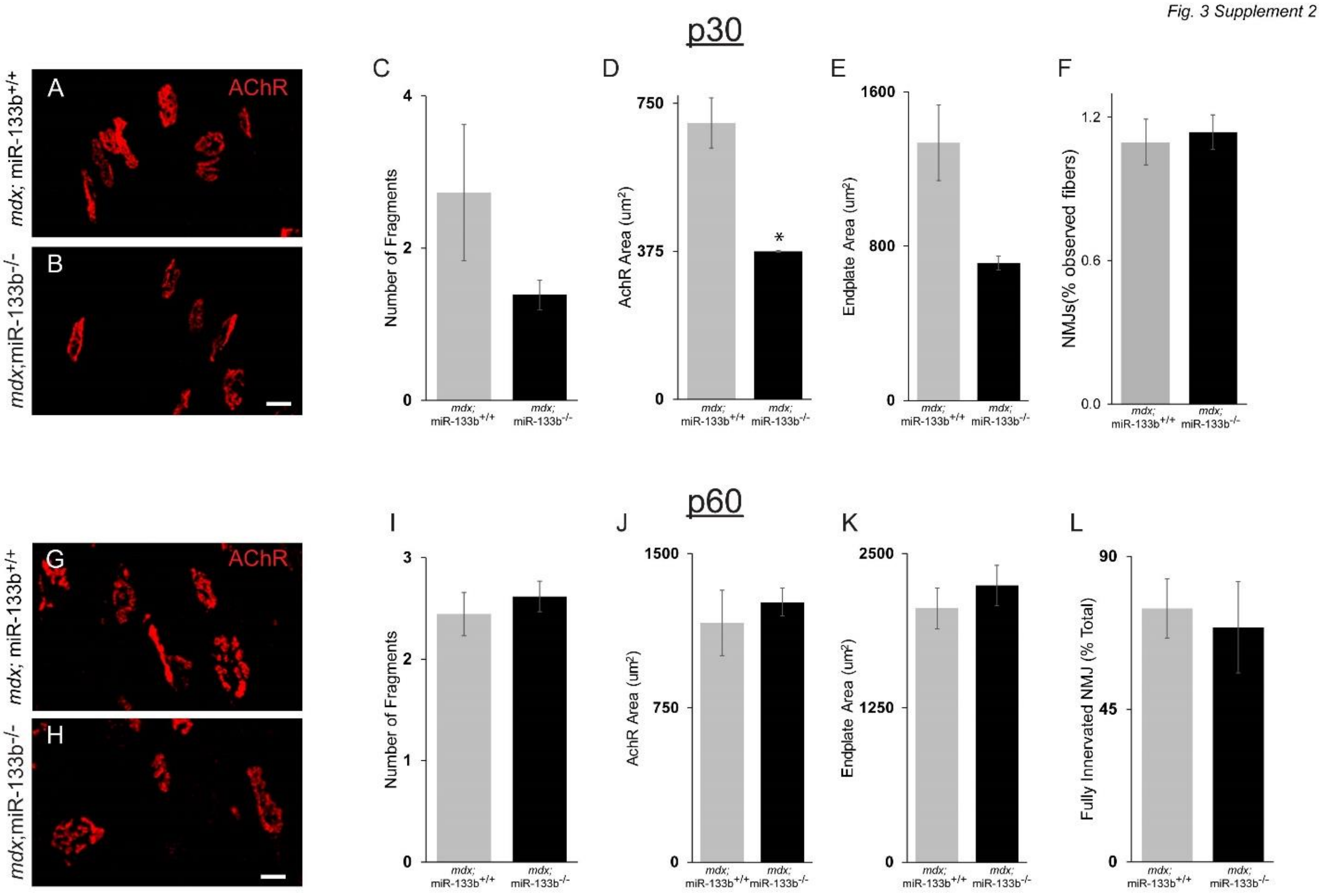


**Supplementary Figure 4.** Deletion of miR-133b affects NMJs in p30 but not p60 *mdx* mice. (A,B) Representative images of AChRs at the NMJ in EDL muscle using fBTX in p30 *mdx*; miR-133b$^{+/+}$ and *mdx*; miR-133b$^{-/-}$ mice. (C-F) Analysis of AChR fragmentation and area, endplate area and NMJ abundance in p30 *mdx*; miR-133b$^{+/+}$ and *mdx*; miR-133b$^{-/-}$ mice. (G,H)) Representative images of AChRs at the NMJ in EDL muscle using fBTX in p60 *mdx*;miR-133b$^{+/+}$ and *mdx*; miR-133b$^{-/-}$ mice. (I-L) Analysis of AChR fragmentation and area, endplate area, and NMJ abundance in p60 *mdx*; miR-133b$^{+/+}$ and *mdx*; miR-133b$^{-/-}$ mice.

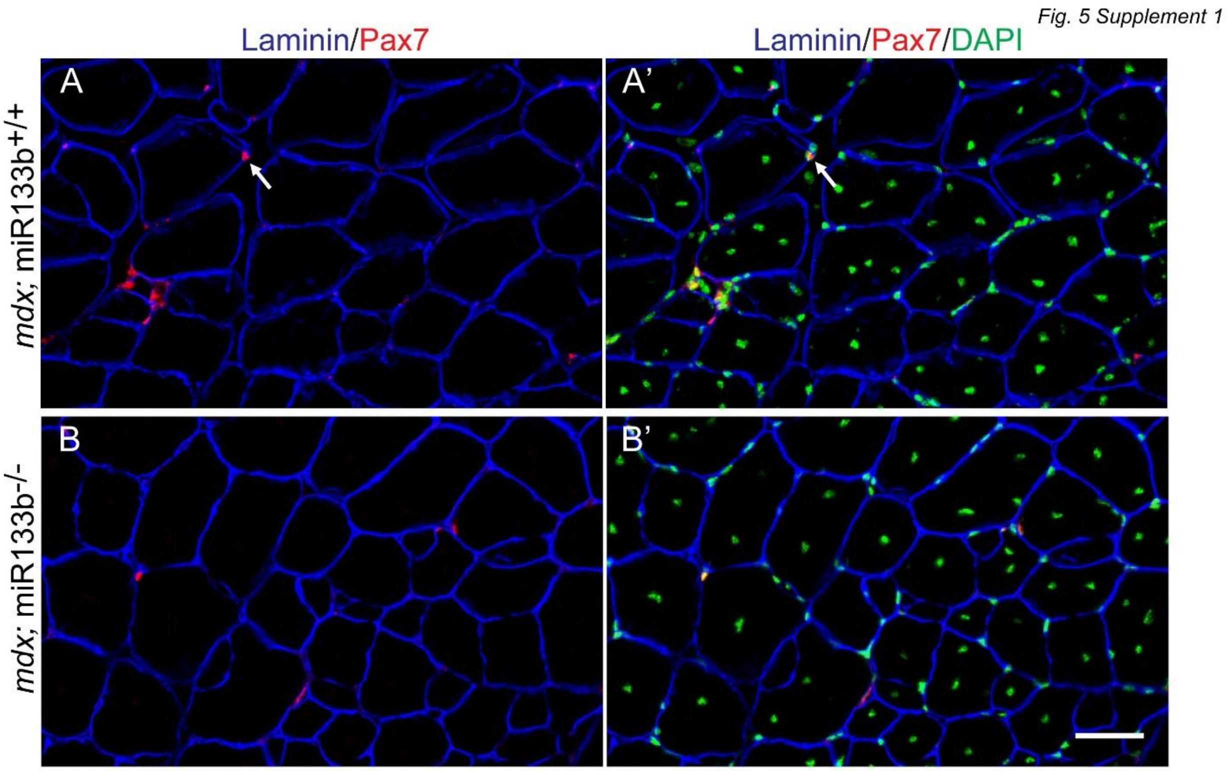


**Supplementary Figure 5.** Representative images of laminin (blue), Pax7 (red) and DAPI (green) immunohistochemistry performed on TA cross sections collected from p90 *mdx*; miR-133b$^{+/+}$ and *mdx*; miR-133b$^{-/-}$ mice. White arrow in panel A indicates an example of Pax7/DAPI double positive nuclei used to identify satellite cells. Scale bar = 50 µm.

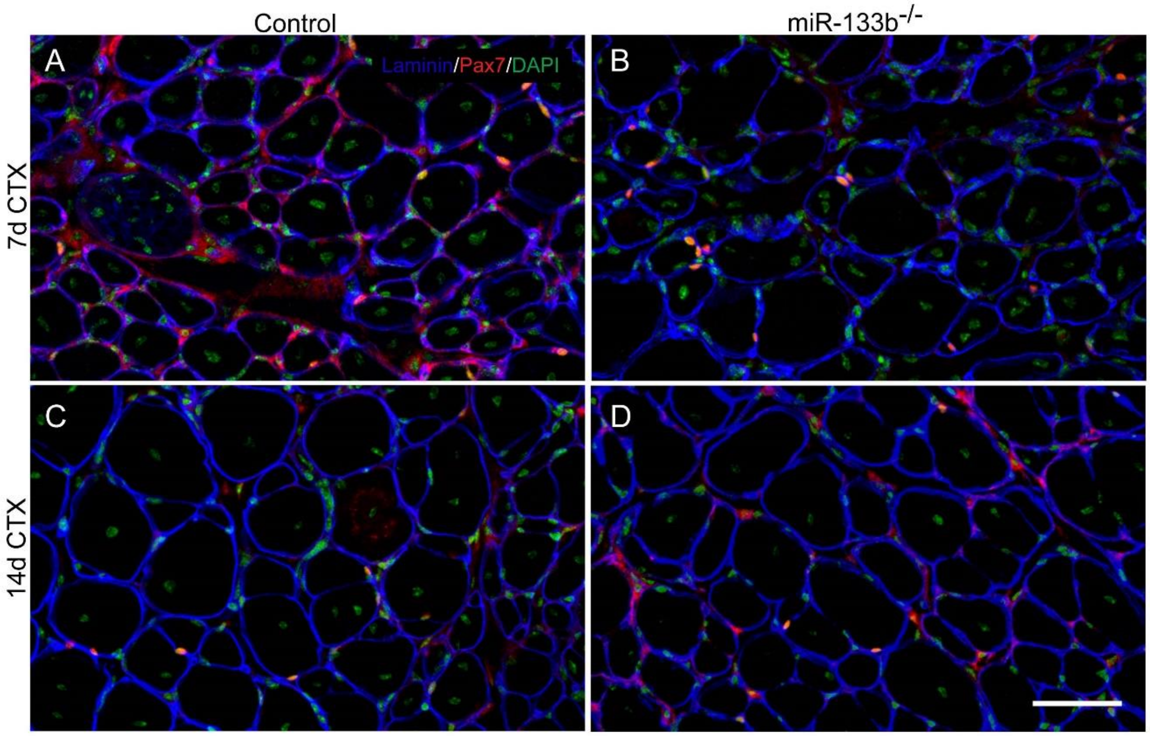


**Supplementary Figure 6.** Representative images of satellite cell abundance in miR-133b$^{+/+}$ and miR-133b$^{-/-}$ TA muscle following cardiotoxin (CTX) administration. Laminin (blue), Pax7 (red) and DAPI (green) immunohistochemistry was performed on TA cross sections collected at (A,B) 7d and (B,C) 14d post-CTX injection. Scale bar = 50 µm.
